# Evidence of widespread balancing selection in a bacterial pangenome

**DOI:** 10.64898/2026.11.28.769857

**Authors:** Cara Conradsen, Edward Feil, Sion Bayliss, Adam Eyre-Walker, Michiel Vos

## Abstract

The vast majority of bacterial genes are only present within a subset of strains in a given species. Such accessory genes may have deleterious or neutral fitness effects and persist only transiently but alternatively might be maintained because they provide a fitness benefit. The adaptive role of accessory genes, and how selection shapes species pangenomes, remains under active debate. Here, we introduce a test that compares the synonymous nucleotide diversity of accessory genes against a neutral coalescent expectation conditioned on accessory gene frequency. Because synonymous variants accumulate approximately clock-like, synonymous diversity acts as a proxy for the time an accessory gene has spent in the pangenome. Discrepancies between observed synonymous diversity and that expected from gene frequency thus provide a signal of differing selective forces. Specifically, cases where synonymous diversity is higher than expected are consistent with long-term balancing selection, while accessory genes with lower than expected diversity indicate directional selection. We apply this test to the accessory genes from 260 representative *Klebsiella pneumoniae* genomes, excluding accessory genes that are likely to have been introduced multiple times. Of 4,071 accessory genes analysed, 1,174 (28.8%) had elevated diversity consistent with balancing selection and 30 (0.7%) had a deficit consistent with directional selection. Among others, we find evidence for balancing selection in genes previously experimentally shown to aid *K. pneumoniae* gut colonisation, as well as in CRISPR-Cas genes, and show directional selection for anti-phage defence system AbiE. Our approach provides the means to disentangle different modes of adaptive evolution that drive the maintenance of accessory genes.

## INTRODUCTION

Bacterial strains of the same species can differ substantially in gene content through the uptake of new genes by horizontal gene transfer (HGT) as well as gene loss (Treangen and Rocha 2011; Vos *et al*. 2015). This phenomenon has led to the concept of the pangenome, in which a relatively small set of core genes is shared by all strains of a species, while an often much larger set of accessory genes (hereafter ‘AGs’) are variably present or absent (Tettelin *et al*. 2005; Domingo-Sananes and McInerney 2021). AGs carry much of the functional variation between strains, including antimicrobial resistance and virulence determinants (e.g., Piper *et al*. 2024), but how they are maintained is still a subject of debate (McInerney *et al*. 2017; Shapiro 2017; Vos and Eyre-Walker 2017; Baumdicker and Kupczok 2023). It is possible that AGs are neutral or deleterious, residing in genomes transiently (Baumdicker *et al*. 2010; Andreani *et al*. 2017), but since AGs may also confer fitness benefits they could also be maintained over longer evolutionary timescales (Lobkovsky *et al*. 2013; Sela *et al*. 2016; McInerney *et al*. 2017). A study using pseudogenes as a neutral reference estimated that ∼20% of AGs across 668 prokaryotic species were adaptive (Douglas and Shapiro 2024). However, a more recent study, investigating whether the coding sequence of accessory genes is constrained, concludes that at least 75% of all AGs in *Escherichia coli* and *Staphylococcus aureus* are adaptive (Eyre-Walker *et al*. 2026).

The discrepancies in the estimates of the proportion of AGs maintained by selection likely reflect conceptual and methodological differences but a further, underappreciated source of variability may be that existing approaches cannot distinguish between different forms of AG adaptive evolution. It is possible that AGs are actively maintained over long timescales through balancing selection, which can be mediated by fluctuating environments (Turner *et al*. 2020), spatial heterogeneity (Liao *et al*. 2021), negative frequency dependent selection (McNally *et al*. 2019; Azarian *et al*. 2020) or even heterozygote advantage when multiple gene copies within an individual are involved (Dewan and Uecker 2025). It is also possible that AGs are under directional selection (Sela *et al*. 2016), resulting in a rapid rise in frequency. However, instead of reaching fixation to become core genes, a more likely scenario is that selective pressures are ephemeral and positive fitness effects turn negative and such variants are short-lived; a recently described process termed ‘adaptive tracking with antagonistic pleiotropy’ (Song *et al*. 2025). This process was recently shown to apply to nucleotide polymorphism in core genes in gut bacteria likely to experience frequently changing conditions (Torrillo and Lieberman 2024).

Novel population genetic frameworks are needed to detect different types of selection that may maintain AGs. Here, we sidestep heterogeneity-based approaches based on different classes of mutations or genes with different expected fitness effects (Vos and Eyre-Walker 2017) to develop a test based on the frequency of AGs as a proxy for the time elapsed since they were first acquired by the species. Our rationale rests on the assumption that AGs under balancing selection persist in the pangenome for longer and should therefore have accrued more synonymous polymorphisms than would neutral AGs that are at the same pangenome frequency (Kimura and Ohta 1973; Bitarello *et al*. 2023). In contrast, AGs under directional selection, as is the case in the adaptive tracking scenario, should have lower synonymous diversity than expected, as they sweep to their observed frequencies. We adapt the age-of-allele test of Blumenstiel *et al*. (2014) developed for transposable element insertions, replacing the within-element molecular clock with a coalescent null model calibrated on core gene synonymous diversity. By comparing observed AG synonymous diversity with a simulated neutral expectation, our test can identify outlier AGs in either tail.

The use of synonymous diversity as a molecular clock for the time since the introduction of an AG would be confounded by multiple introductions; when different alleles of the same AG are taken up into the species pangenome, nucleotide diversity will be inflated. To account for this effect, we implement two distinct tests. First, we exclude AGs that represent statistical outliers with respect to a high synonymous diversity. Second, because core gene order is highly conserved (Rocha 2006; Rocha 2008) and AGs can be assigned chromosomal context, we remove non-syntenic AGs that might have arisen through multiple independent uptake events. We note this dual approach is conservative, as high nucleotide diversity might also arise due to mutation and homologous recombination within the focal species, and non-synteny could also arise due to intragenomic rearrangements, or recombination within the pangenome rather than via multiple introductions (Arnold *et al*. 2022).

We apply our test to a temporally and geographically constrained *Klebsiella pneumoniae* SpARK dataset (Thorpe *et al*. 2022). Of the 4,071 singly introduced AGs, 29.6% depart from neutral expectation, overwhelmingly in the direction of high synonymous diversity. This indicates long-term maintenance by balancing selection predominates over transient adaptive tracking, although directional selection may be harder to detect. We note detection of two distinct signatures of balancing selection: genes that are maintained within a subset of strains, and cases in which both the gene and divergent alleles within the gene are maintained, as exemplified by candidates including O-antigen biosynthesis genes, CRISPR-Cas loci, and type VI secretion components. Candidates for directional selection are rarer but include the AbiE phage-defence operon, in which both toxin and antitoxin carry fewer synonymous variants than expected. By distinguishing between balancing and directional selection, our approach provides a means of determining how selection maintains accessory gene variation, a central and unresolved question in bacterial evolution.

## RESULTS

### Klebsiella pneumoniae pangenome

We describe a test for selection on accessory genes based on comparing the observed number of segregating synonymous sites with neutral expectations derived from gene frequency. We use the SpARK One Health collection (SpARK; Thorpe *et al*. 2022) of clinical, agricultural, and environmental *K. pneumoniae* isolates as our test dataset (Supplementary Fig. 1a). This genome collection was collected over a 17-month period in 2017-2018 in and around the city of Pavia, Northern Italy. The use of a geographically and temporally constrained sample minimises potential confounders arising through isolation by distance or evolution through time. We selected 260 closed genomes, assembled from combined short- and long-read data, distributed across 108 sequence types (STs), of which the most prevalent were ST307 (19.6%), ST512 (6.2%) and ST661 (6.2%) (Supplementary Fig. 1b). A total of 18,486 genes were identified using PIRATE (Bayliss *et al*. 2019; Supplementary Fig. 1c). These follow the typical pangenome asymmetric U-shape gene frequency distribution (Baumdicker *et al*. 2012; Collins and Higgs 2012; Haegeman and Weitz 2012; Lobkovsky *et al*. 2013), with the bulk of genes being cloud (present in ≤ 15% of genomes; 72% of pangenome genes) and core (present in ≥99% of genomes; 22%; Supplementary Fig. 1d).

### Detecting multiply introduced accessory genes

Our test rests on the assumption that AGs accumulate synonymous diversity over time after a single introduction into the pangenome. Yet in *K. pneumoniae*, high HGT rates mean the same AG could potentially be introduced repeatedly (Navon-Venezia *et al*. 2017; Wyres *et al*. 2020; Haudiquet *et al*. 2021). When multiple introductions carry divergent alleles, synonymous diversity will be inflated relative to that from a single introduction. We used two filter steps to control for this effect. After removing paralogs and singletons, which we cannot apply our analysis to (Table 1), we filtered out 611 AGs with substantial synonymous diversity estimates, defined as exceeding the median by more than three interquartile ranges (median *π_S_* = 4.76 × 10⁻³; threshold *π_S_* = 0.10; Fig. 1a). Second, we implemented a filter based on synteny, reasoning that an AG found between the same pair of core genes is likely to be the product of a single introduction. We followed Molari *et al*. (2025) in using PanGraph (Noll *et al*. 2023) to identify collinear core genes that we group into core regions (‘core-blocks’), in our aligned genomes, in order to account for large scale inversions and translocations (Fig. 1b). AGs were then mapped onto these core-blocks between flanking core genes (Oliveira *et al*. 2017; Molari *et al*. 2025). Thirty core-blocks were identified, with conserved gene order across 238 of 260 isolates (91.54%). Of the 22 genomes in which block order was not conserved, the number of blocks impacted by rearrangements ranged from 1 to 19 (median = 4; Supplementary Fig. 2). The core-blocks establish a robust reference for tracking AG location and synteny without the need to rely on a reference alignment. In total, 4,071 AGs were consistently localised between the same pair of flanking core genes within the same core-blocks (Table 1).

**Figure 1.**
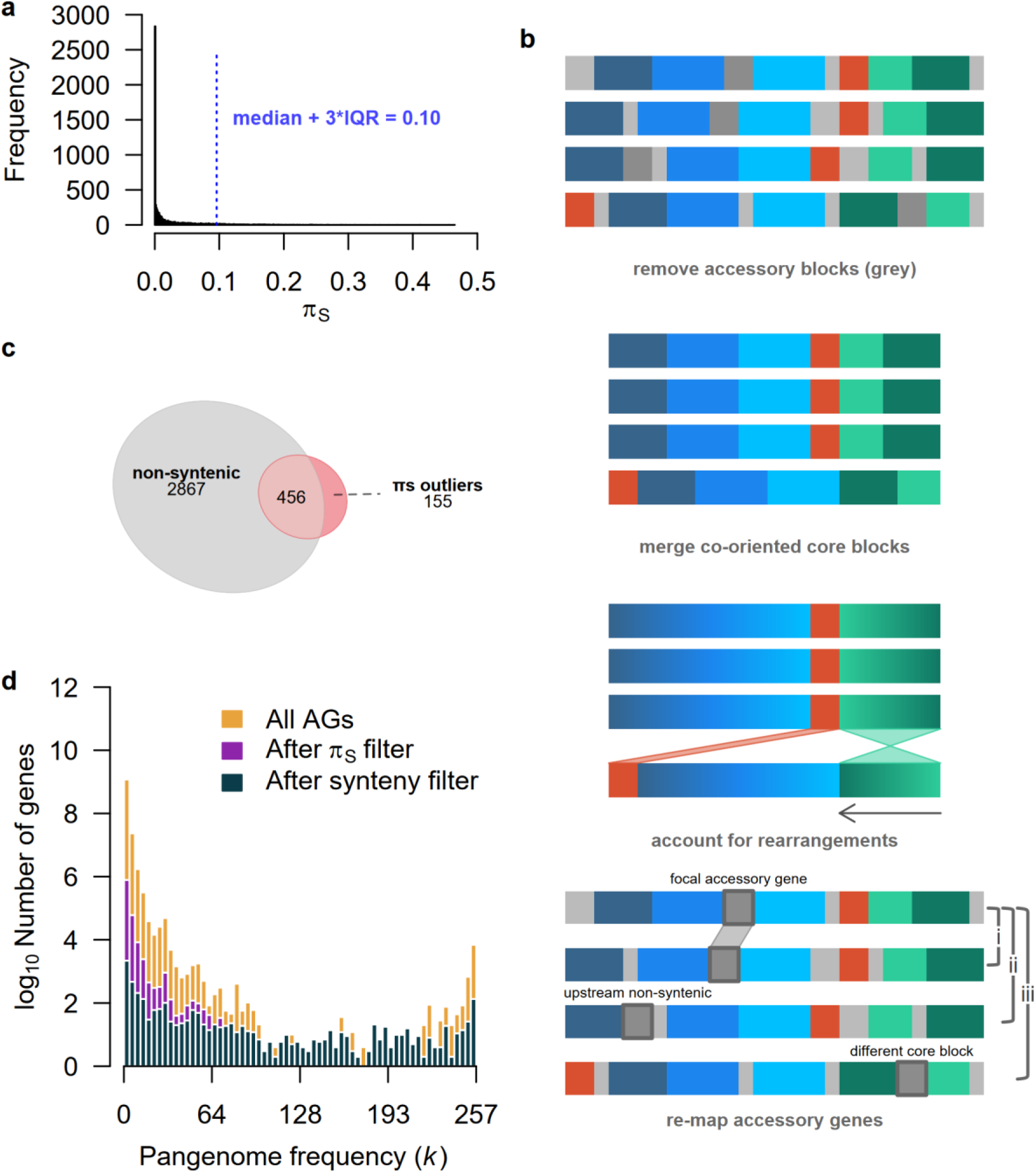
Filtering out multiply introduced accessory genes (AGs) using nucleotide diversity and synteny criteria. (a) Distribution of synonymous diversity (*π_S_*) and the diversity threshold (median + 3 × interquartile range, blue dashed vertical bar). (b) Schematic of accessory gene localisation using *Pangraph* synteny alignments. Accessory (grey genes) and core (coloured genes) genomic regions are identified across genome alignments (top panel). Accessory genes are removed and collinear core genes are merged into ’core-blocks’ (similar hues: blue, teal and orange), accounting for translocations and inversions (middle panel). AGs are then mapped to core-blocks based on their flanking core context (bottom panel). Two copies of an AG are considered syntenic, and hence representing a single introduction, when found between the same pair of flanking core genes within a core-block (e.g., focal AG and AG i, both within the blue segment). AGs are considered non-syntenic when located between different core genes within the same core-block (focal AG vs AG ii) or in different core-blocks entirely (focal AG vs AG iii). (c) Venn diagram summarising AGs with multiple inferred introductions identified across the two filtering processes. Numbers represent genes counts within each individual set (circle fraction). (d) Stacked histogram showing the number of AGs retained (y - axis, log_10_ scale) at different pangenome frequencies (*k*, the number of genomes carrying the AG, binned in intervals of 4). Bars are shaded from lightest to darkest to represent the proportion remaining after each sequential filtering step for singly acquired AGs: starting with all AGs (yellow); after removing *π_S_* outliers (magenta); and, after removing non-syntenic AGs (teal-navy).

**Table 1.** Stepwise filtering for singly introduced accessory genes (AGs) . Summary of gene counts across sequential filtering steps used to identify AGs for the selection analysis (see Methods).

| **Analysis** | **Method** | **Gene Category** | **Gene Count** | **AGs after filter step** |
|---|---|---|---|---|
| Establishing pangenome | PIRATE | Total genes | 18,486 |  |
|  |  | Paralogs | 614 | 17,872 |
|  |  | Core | 3,851 | 14,021 |
|  |  | Single-genome accessory genes | 6,472 | 7,549 |
| Removal of genes associated with multiple introductions | High π<sub>S</sub> filter | π<sub>S</sub> outliers | 611 | 6,938 |
|  | PanGraph synteny analysis | Non-syntenic | 2,867 | 4,071 |

The synteny and high *π_S_* filters were not independent. AGs at inconsistent genomic positions had an 8.8-fold higher median *π_S_* than those confined to a single syntenic locus (0.016 versus 0.0018; *n* = 3,323 and 4,226; Wilcox rank-sum test, *P* <0.001, before *π_S_* filtering), and AGs excluded for high *π_S_* were more often non-syntenic than those retained (75% of 611 versus 41% of 6,938, Fig. 1c; Fisher’s exact test, OR = 4.18, *p* < 0.001). Consequently, 456 AGs (13% of the 3,478 excluded by either filter; Fig. 1c) were removed by both filters. This overlap was expected as both filters were designed to catch multiple introductions. Each filter mostly excluded low frequency AGs (Fig. 1d), which are known to be dominated by mobile genes (Gamblin *et al*. 2025).

### Identification of accessory genes under selection

Balancing selection predicts an excess of synonymous polymorphisms in AGs relative to their frequency in the population, whereas directional selection predicts a deficit. To detect AGs deviating from the neutral expectation, we compared the observed number of synonymous segregating sites in each singly introduced AG to coalescent-derived expectations across all gene frequency classes (ranging from 2 to 257 genomes). The *π_S_* estimate was lower than Watterson’s estimator (0.0121 versus 0.0195, *π_S_*/*θ*_W_ = 0.62), consistent with an excess of rare variants (i.e. a negative Tajima’s D; Tajima 1989) and suggesting the population is under expansion. We therefore conducted our coalescent simulations under a variety of exponential population size expansion models varying the expansion parameter, α; we found that an expansion rate of 2.8 and a *θ* value of 0.031 matched the observed values of *π_S_*/*θ*_W_ and *π_S_* (Supplementary Fig. 4a-c). When accounting for population size expansion, the 95% neutral interval moved downward and narrowed in range, suggesting that fewer low diversity outliers could be detected (but with high diversity outliers unaffected) (Fig. 2a). Inference of directional selection is therefore sensitive to the assumed demography, whereas inference of balancing selection is robust to it.

**Figure 2.**
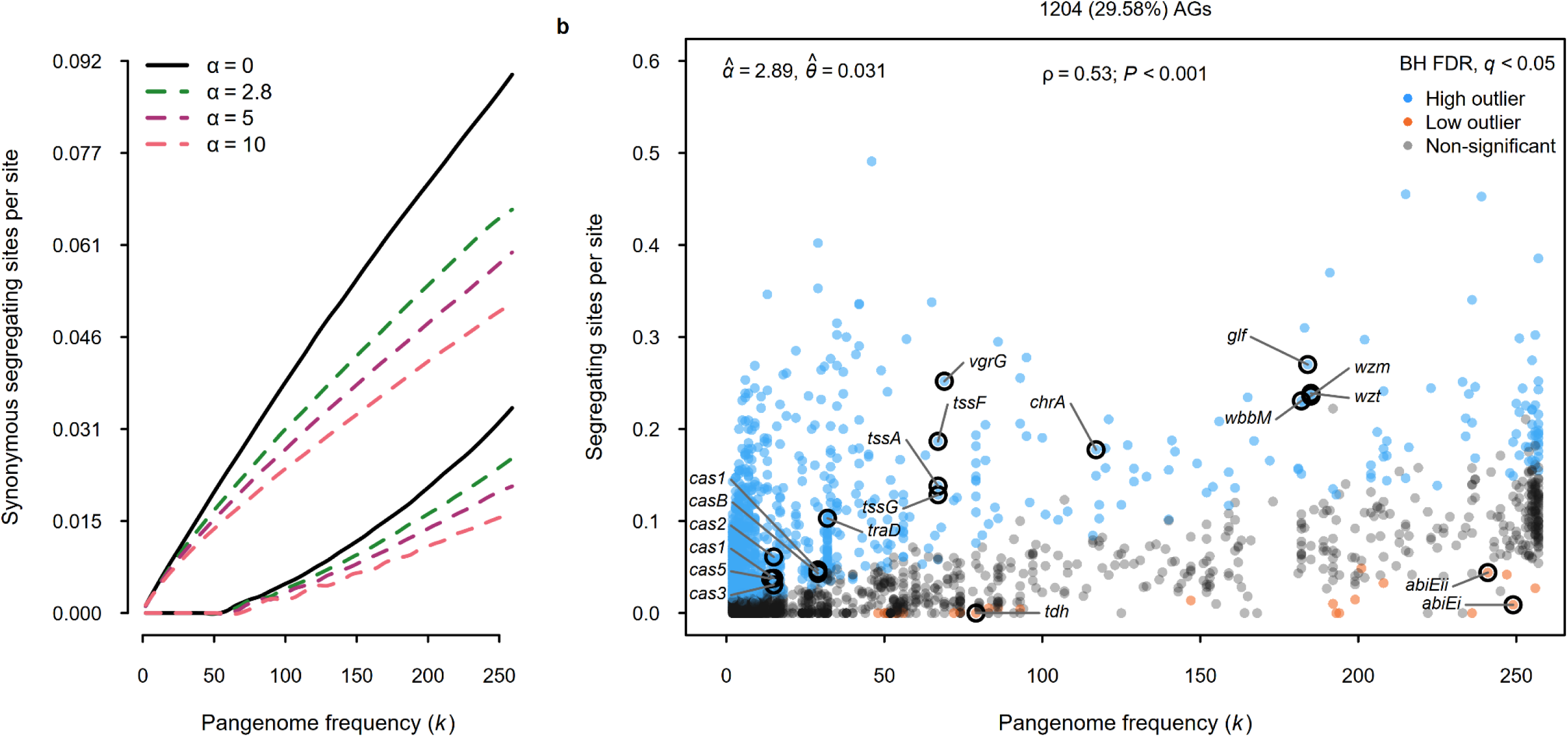
Neutral expectation for synonymous diversity and identification of accessory gene (AG) outliers. (a) The expected number of synonymous segregating sites as a function of pangenome frequency (*k*) under four expansion scenarios (growth rate, α = 0, 2.8, 5 and 10). Curves are drawn at fixed θ, taken as the observed core-genome *π_S_*, to highlight the effect of demography on the shape of the neutral interval. For each scenario, the upper and lower curves delimit the 95% interval. The solid black line shows this interval under a constant size population (α = 0); with dashed lines for expanding populations. Expectations were derived from 500,000 coalescent genealogies simulated in *msprime* (Baumdicker *et al*. 2022). (b) Number of synonymous segregating sites per site for each AG plotted against pangenome frequency (*k*), under an expansion model where α^ = 2.89 and θ^ = 0.031. Model parameters, α^ and θ^ were estimated sequentially, since a higher interpolated growth rate required a higher θ to reproduce the observed core-genome *π_S_* (Supplementary Fig. 4). A Benjamini-Hochberg false discovery rate correction was applied across all AGs, where high (blue) and low (orange) outliers have adjusted *q* < 0.05 above and below the neutral expectation, respectively. Numbers above the panel indicate the count and percentage of outlier AGs. Spearman’s rank correlation (*ρ*) between observed synonymous segregating sites and pangenome frequency across all AGs is shown in the top centre. Non-significant AGs are shown in grey. Labelled genes belong to multi-component systems for which two or more members were independently recovered as outliers, together with named transporters and enzymes with assignable function. Full outlier list in Supplementary Table 1.

As expected, synonymous diversity co-varies with frequency across all AGs (Spearman’s ρ = 0.53, *P* < 0.001; Fig. 2b); however, when comparing the observed synonymous diversity against neutral expectations 1,204 AGs (29.58%) departed from the null under an FDR of 5% (Fig. 2b, Supplementary Table 1). Most outliers (*N* = 1,174, 97.5%) had more synonymous segregating sites than expected under neutrality, consistent with balancing selection (Fig. 2b). Accounting for these outliers under a neutral model would require a mutation rate several-fold higher than the fitted estimate of 0.031, which was itself constrained by the core genome (θ^ = 0.0308 [0.0308, 0.0309]; Supplementary Fig. 4b, c), indicating that they cannot be explained by underestimation of *θ*. While 85.9% of high diversity outliers occurred at low frequency (in less than 50 isolates; Fig. 2b), so do 81.5% of non-outlier AGs. The apparent concentration at low frequency therefore simply reflects the underlying frequency distribution of all AGs.

Only 30 AGs had fewer segregating synonymous sites than expected, consistent with directional selection, and these were concentrated at intermediate to high frequency (median *k* = 81, lowest *k* = 48; Fig 2b). It is possible the small number of low diversity outliers reflects limited power of the test rather than a rarity of sweeps. A deficit can only reach significance when the neutral expectation is large enough that observing few or no segregating sites is improbable at the realised Benjamini-Hochberg threshold (*P* = 0.0146), this requires the expected number of synonymous segregating sites to exceed 4.9 (Supplementary Fig. 5a). As the number of expected sites scales with synonymous length and pangenome frequency, a median gene synonymous length (*L*_S_ = 237) first becomes testable at *k* = 63 (Supplementary Fig. 5b), where only 570 of 4,071 AGs (14%) could return a low outlier call. Detecting sweeps at lower frequencies would require substantially more genomes.

### Characteristics of accessory genes departing from neutral expectation

Outlier AGs were distributed across 22 out of 26 COG categories (Fig. 3a). After FDR correction, balancing selection candidates were over-represented in the mobilome (X; odds ratio: 1.58, *q* < 0.01, Fig. 3a and b), as previously noted (Corander *et al*. 2017; White *et al*. 2022). Another category where high-diversity outliers were overrepresented was replication, recombination and repair machinery (L; 1.73, *q* < 0.01), while the category cell wall, membrane and envelope biogenesis (M; 0.64, *q* < 0.05) was depleted (Fig. 3b). When enrichment comparisons were made among high-diversity outliers stratified by AG type, two categories that had previously been nominal reached significance among cloud AGs: carbohydrate transport and metabolism was depleted (G; 0.61, *q* < 0.05) and extracellular structures enriched (W; 6.29, *q* < 0.01), the strongest single enrichment in the dataset. The latter was masked in the pooled comparison (OR 2.08, *q* = 0.18) by an opposing but non-significant trend among shell AGs (OR 0.50, from two genes). No category reached significance in the shell or soft-core strata, which contained only 118 and 40 outliers, respectively. Among low-diversity outliers, no COG category survived correction. Three cellular processes and signalling categories showed nominal signal: signal transduction mechanisms (T; odds ratio 4.15, *P* = 0.048), intracellular trafficking, secretion and vesicular transport (U; 4.49, *P* = 0.040) and defence mechanisms (V; 5.26, *P* = 0.006).

**Figure 3.**
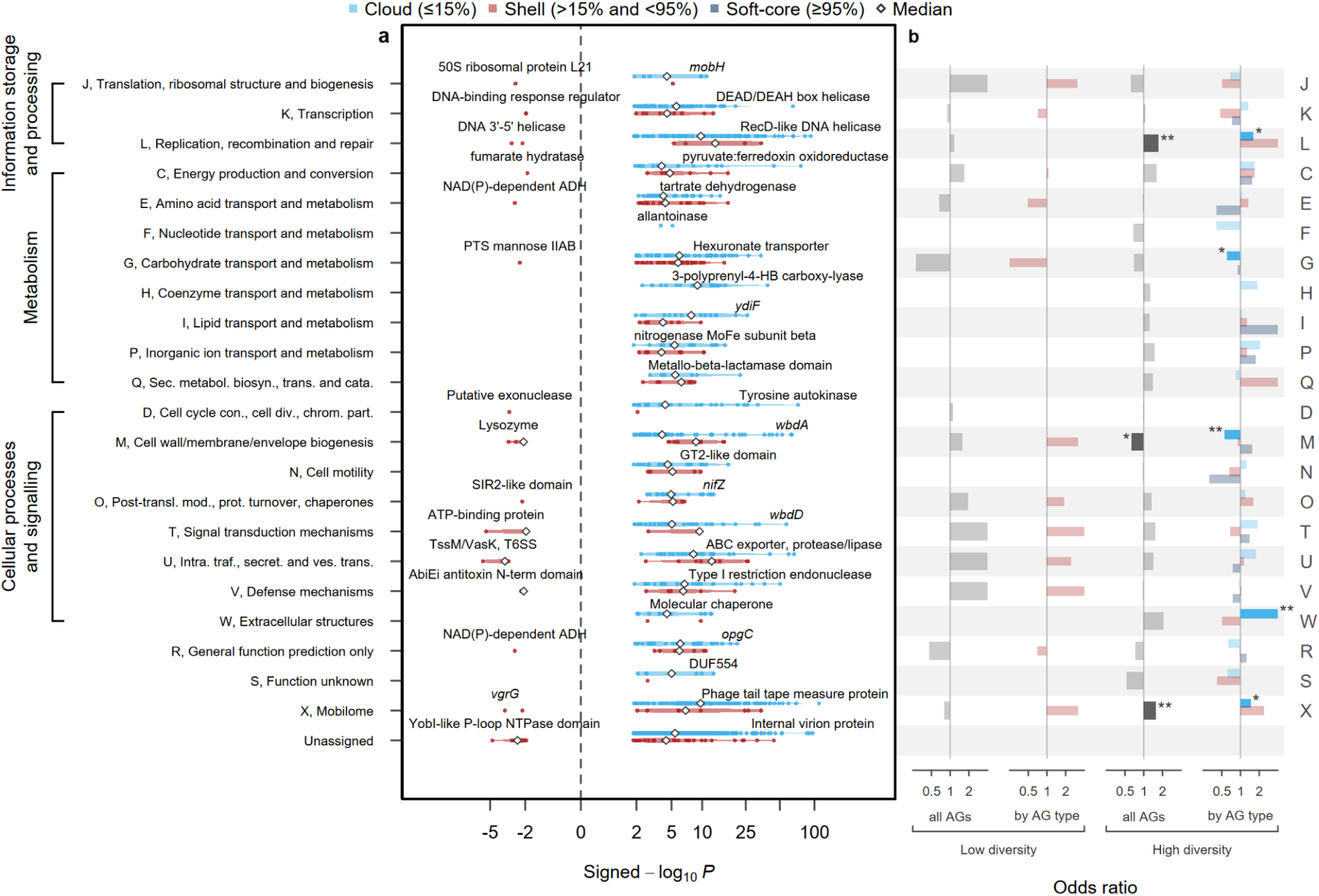
Selection signatures in *Klebsiella pneumoniae* accessory genes (AGs) by COG functional category, split by pangenome frequency, with functional enrichment. **(a)** Distribution of departure from neutrality among outliers. Signed −log₁₀(p) values for AG outliers (*N* = 1,204: 1,174 with excess diversity and 30 with reduced synonymous diversity) were computed per gene from a two-sided test against the neutral distribution of synonymous segregating sites, a Poisson mixture over simulated genealogies (see Methods), for each gene’s frequency and synonymous length. The sign indicates direction relative to neutral expectation (vertical dashed line, signed −log₁₀(*p*) = 0); positive values indicate excess synonymous diversity (consistent with long-term maintenance by balancing selection); negative values indicate a deficit (consistent with a shorter than expected presence in the pangenome for a given frequency due to a selective sweep). The x-axis is drawn on a signed log-modulus scale, sign(x)·log₁₀(1+|x|), with ticks in the original units. Within each COG category, genes are divided into cloud (≤15%), shell (>15% and <95%) and soft core (≥ 95%). Violins show the density of genes, white diamonds represent the median, each drawn separately for high and low sides of zero and only where there were at least three genes in the relevant frequency class. Genes with extreme values in their category are labelled. COG categories are grouped on the left by higher-level functional classes; some genes (*N* = 158) were assigned to more than one COG category and are plotted in each. Abbreviated categories: D, cell cycle control, cell division, chromosome partitioning; O, post-translational modification, protein turnover, chaperones; U, intracellular trafficking, secretion and vesicular transport; Q, secondary metabolites biosynthesis, transport and catabolism. (b) Functional enrichment odds ratios are from two-sided Fisher’s exact tests of COG membership. Each outlier category (high vs low) was tested against non-outliers separately. The first column of each pair pools all AGs (grey bars); the second repeats the test within each frequency class, coloured and vertically offset as in a. Darker bars and asterisks mark categories passing Benjamini–Hochberg correction within that comparison (* *q* < 0.05, ** *q* < 0.01, *** *q* < 0.001). Categories with no bar were not returned for that comparison.

Having used synonymous variation to measure genealogical depth, we next looked at variation in outlier genes at the protein level. We quantified amino acid sequence variation using protein allelic diversity (*H*), calculated as Nei (1978)’s unbiased gene diversity over the frequencies of distinct translated sequences, so that *H* is the probability that two genomes carrying the AG encode different protein variants. We expected low synonymous diversity outliers to have low *H* because recent sweeps leave insufficient time for protein variants to accumulate. However, high synonymous diversity outliers could display a range of *H* values. High synonymous diversity outliers with low *H* point to selection on the level of the AG itself, where AG sequence is conserved and it is the presence or absence of the AG *per se* that is favoured, whereas high synonymous diversity outliers with high *H* could indicate that the alleles within the AG are subject to balancing selection.

As expected, low outliers had little protein variation (median *H* 0.09, interquartile range, IQR: 0 - 0.24, Fig. 4a). In contrast, high outliers spanned the full range of *H* (median 0.71, IQR 0.49 - 0.93). *H* increased monotonically with synonymous diversity; the association was strongest among non-outlier AGs (Spearman’s ρ = 0.52, *P* < 0.001) and low outliers (ρ = 0.45, *P* = 0.012), and weakest among high outliers (ρ = 0.32, *P* < 0.001). This weaker association could suggest that variation in *H* among high outliers is driven by selection for allelic richness rather than genealogical depth. However, because the outlier classes are defined by their largely non-overlapping synonymous diversity, a common model of *H* on synonymous diversity cannot be fitted to test this. We found no significant enrichment after correction of co-occurring AGs among outliers (Supplemental Information, Supplementary Fig. 6), and limited information on strain origin precluded association of outlier AGs with specific environments (Supplemental Information).

**Figure 4.**
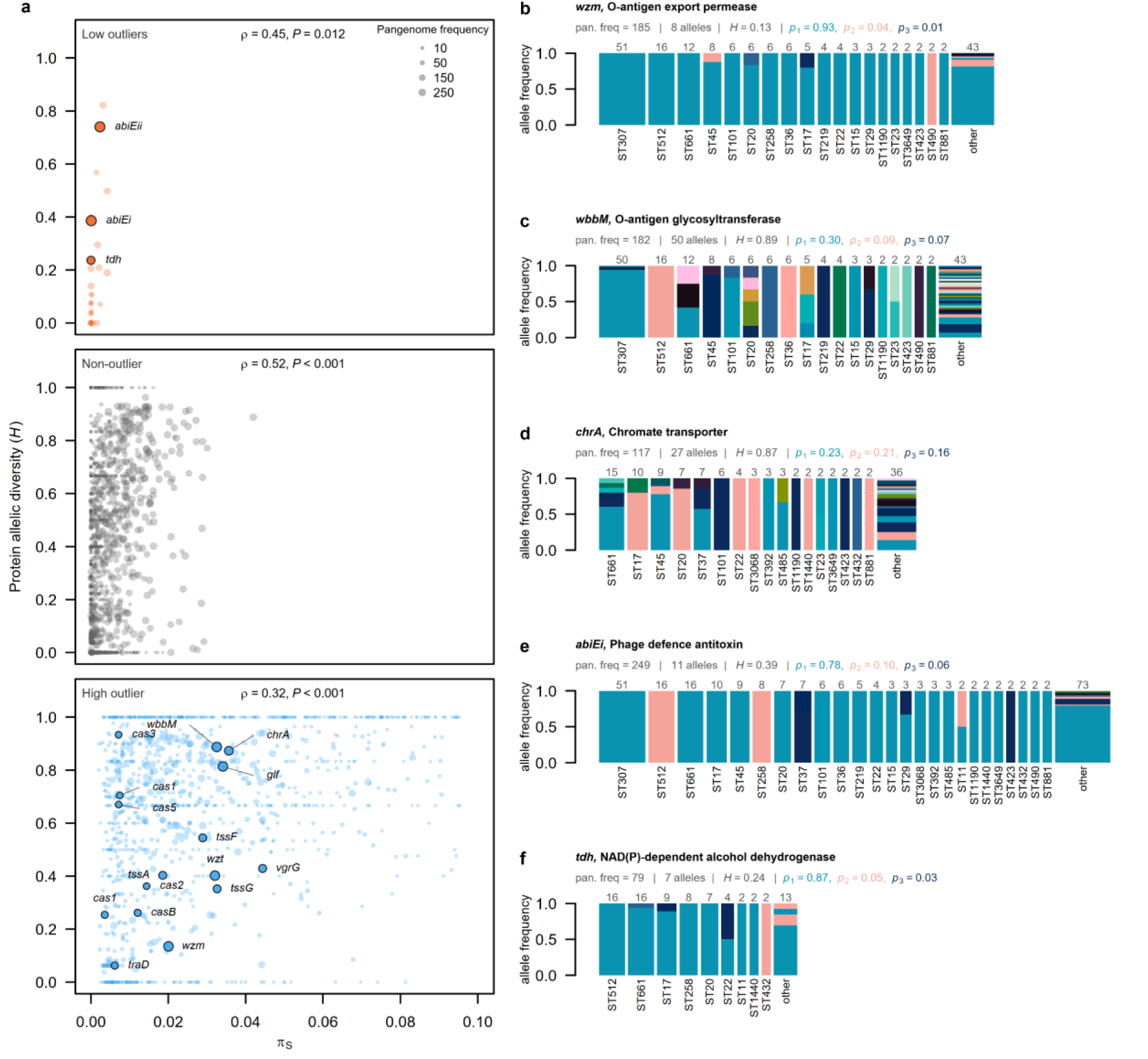
Protein diversity of accessory genes. (a) Nei (1978)’s unbiased protein diversity, *H*, against synonymous nucleotide diversity, *π_S_*, for AGs that were low outliers (top), non-outliers (middle) and high outliers (bottom), gene labels and point colours as in Fig 2b. *H* was calculated classifying protein sequences as distinct alleles; where Spearman’s rank correlation, *ρ*, between *H* and *π_S_* is given at the top of each panel. Point size scales to pangenome frequency (*k*). (b - f) Protein allele frequencies by sequence type (ST) for the AGs *wzm*, *wbbM, chrA, abiEi* and *tdh*. Bar width is proportional to the square root of the number of genomes carrying the gene in each ST, on a scale common to all panels; numbers above bars give these counts. STs with a single carrier of the gene are pooled as “other”. Alleles are ranked and coloured by overall frequency. The three commonest (*p_1_*, *p_2_*, *p_3_*) are shown in teal, pink and navy in every panel, and their frequencies are given in the same colours in each panel header, alongside *k*, the number of alleles and *H*.

### Examples of accessory genes under balancing and directional selection

It is not possible to review all 1000+ AGs identified to be under selection (Supplementary Table 1), but below we highlight a selected range to illustrate the power of the test to identify both ‘usual suspect’ genes and genes hitherto not known to be under selection. Antigens are prime candidates for host-mediated balancing selection (Brisson and Dykhuizen 2004). All six high diversity outlier genes at the focal O-antigen locus (*k* = 182–185) lie within the same co-occurrence cluster (cluster 27; Supplementary Table 1, Supplementary Fig. 6). Two of them, *wzm* (46 synonymous segregating sites against 14.4 expected; *q* < 0.001) and *wbbM* (116 against 36.4; *q* < 0.001), show contrasting patterns of molecular variation: although they are of near-identical frequency (*k* = 185 and 182; Fig. 2b) and lie in between the same flanking core genes, they differ almost sevenfold in protein allelic diversity (*H* = 0.13 versus 0.89; Supplementary Table 1; Fig. 4b, c), a difference that is not explained by gene length difference (Supplementary Fig. 7). One *wzm* protein variant is carried by 93% of 185 genomes across 54 STs, indicating that selection acts to maintain the presence of one main version of this transporter gene. The *wbbM* gene shows a very different pattern, having 50 protein variants, the commonest of which has a frequency of 30% spanning only six STs, which could point at additional selection on the level of alleles to change O-antigen structure in response to parasite-host coevolution, although we do not have evidence for this process.

We found significantly elevated synonymous diversity in six *cas* genes distributed across two type I-E CRISPR-Cas loci among cloud genes (*cse2/casB* and *cas1e* at the first locus, *k* = 29, *q* = 0.018 and *q* = 0.002, respectively; *cas3, cas1e, cas5e* and *cas2e* at the second locus, *k* = 14–15, *q* < 0.001; Fig. 2b, Supplementary Table 1). CRISPR-Cas is an adaptive immunity system engaged in antagonistic coevolution with MGEs, making it a candidate for negative frequency-dependent balancing selection. Negative frequency dependent selection leading to spacer acquisition has been demonstrated experimentally (Common *et al*. 2019), but selection on *cas* genes themselves has rarely been examined. Takeuchi *et al*. (2012) found evidence for purifying selection in CRISPR-Cas genes using the dN/dS test. As is the case for the *wzm* transporter above, our evidence for balancing selection based on long-term allelic persistence does not conflict with this result: we find that within species, *π_N_*/*π_S_*< 1 indicating that the protein sequence is conserved but we show that these genes have higher levels of synonymous diversity suggesting that balancing selection acts on the presence/absence of the genes themselves.

The chromate transporter *chrA* (*k* = 117; 60 synonymous polymorphisms against 14 expected; Supplementary Table 1; Fig. 2b, Fig. 4d) carries 27 protein variants, the three commonest each at 16–23% frequency and spanning 8–11 STs. Balancing selection on *chrA* could potentially be linked to observed elevated chromium concentrations in the Po Valley due to geological weathering (Amorosi *et al*. 2010; Amorosi *et al*. 2014) and potential anthropogenic pollution. To our knowledge this is the first finding of selection on a chromate resistance gene in natural populations, which complements functional evidence (Branco *et al*. 2008). No mobile elements were detected at this locus in any genomes, despite known plasmid or transposon association of chromate resistance genes (Amorosi *et al*. 2010). Several other resistance-associated AGs were identified as balancing selection candidates, including both subunits of a predicted EmrAB-type multidrug efflux pump (HlyD-family periplasmic adaptor; *k* = 142; 43 segregating synonymous sites vs. 13.4 expected; and, DHA2-family MFS transporter; *k* = 141; 76 vs. 22.0), as well as the urease accessory protein *ureD* which plays a role in virulence (*k* = 13; 11 vs. 0.78). We also uncovered multiple HGT-related outlier AGs, including chromosomally located plasmid-derived conjugation and mobilisation genes (*traD*, *k =* 32, 64 vs. 5.6; *traC*; and *trbL*; Supplementary Table 1) and the replication protein RepA of chromosomally integrated elements.

Our evidence for balancing selection for a variety of genes is in part corroborated by a *K. pneumoniae* transposon mutagenesis screen using a murine model of GI colonisation with an intact microbiota which focused on genes encoding membrane-associated proteins (Bray *et al*. 2025). We find evidence for selection on *tssA*, *tssF*, *tssG*, *vgrG* (five gene families) and *vasK* genes (Fig. 2b), which are components of the bacterial type VI secretion system (T6SS) which acts like a contractile molecular spear to deliver effector proteins into neighbouring cells, while the transposon screen identified T6SS genes *tssB*, *tssG*, *tssJ, tssK* and *clpV* as aiding gut colonisation by facilitating competition with *Betaproteobacteria* species (Supplementary Table 1). Our test found evidence for balancing selection in cytochrome c biogenesis ATP-binding export protein *ccmA* and Type 1 fimbrial protein *fimA* alongside 27 other (putative) fimbrium-associated genes (Supplementary Table 1), which the transposon screen uncovered as genes facilitating gut colonisation.

Candidates for directional selection (i.e., those with little synonymous diversity) include the AbiE phage-defence operon. The antitoxin *abiEi* is among the most extreme deficit outlier in the dataset, carrying 2 synonymous segregating sites against 22.9 expected (*k* = 249), and the cognate toxin *abiEii* is also a low diversity outlier (11 against 24.7, *k* = 241; Supplementary Table 1; Fig. 2b, Fig. 4e). A single AbiEi protein variant is carried by 78% of the 249 genomes and shared across 80 STs, consistent with a recent sweep, possibly driven by phage-mediated selection pressure. The NAD(P)-dependent alcohol dehydrogenase *tdh* is also notable. It had no synonymous polymorphism (vs. 8.3 expected segregating sites), yet occurred in 79 genomes across 24 STs, was absent from the dominant ST307 clone and was located outside any mobile element (Supplementary Table 1; Fig. 2b, Fig. 4f). Its seven protein alleles therefore sit on a single synonymous background, the commonest of which is shared by 87% of genomes carrying this AG across 19 STs. *tdh* co-occurs with a complete sugar catabolic operon whose other genes (*ydjI*, *ydjK*, *ydjL* and *glpR*) also have few synonymous sites (1-2 vs. ∼ 8), although not significantly so. This, together with its intermediate frequency, is suggestive of an adaptive tracking signature where the AG is mid-trajectory during a selective sweep.

## DISCUSSION

Debate over the evolution of the accessory genome has centred on the proportion of genes under selection, with little attention to the modes of selection acting upon them. This in large part reflects the lack of suitable population genetic frameworks to distinguish balancing selection from directional selection. Here, we develop a test of selection on the genic level based on the deviation of empirical AG frequency and synonymous diversity against a coalescent-derived neutral expectation to identify AGs that are under balancing or directional selection and apply it to a curated bacterial pangenome dataset. We find evidence of widespread long-term maintenance of AGs, including in CRISPR-Cas and other genes for which there was no previous evidence of adaptive evolution based on traditional d_N_/d_S_ approaches.

Our estimate that 29.6% of AGs are under some form of adaptive evolution sits between the ∼20% of AGs inferred using a pseudogene reference (Douglas and Shapiro 2024) and a lower bound of 75% inferred from *π_N_*/*π_S_* by Eyre-Walker *et al*. (2026) and comes with two main caveats. First, our method cannot interrogate singleton AGs, and we filtered out AGs that displayed hallmarks of possible repeat introduction, leaving only 4,071 out of 14,021 (29.0%) AGs available for testing. Second, within this set, our test is unequally powered across genes, with 86% of AGs below the threshold where we could detect low diversity outliers (Supplementary Fig. 5) and our estimate thus represents a lower bound of adaptive evolution.

Balancing selection outliers were predominantly found at low frequencies (i.e. cloud genes), which reflects the underlying AG distribution (and contrasts with the intermediate frequencies at which it is often sought; Corander *et al*. 2017). We found that AGs interpreted to be under significant balancing selection ranged widely in protein diversity, which potentially could be due to balancing selection on the level of alleles in addition to that of AG carriage; further work is needed to test whether this is indeed the case. We found limited evidence for directional selection where AGs reach their observed frequency on short timescales and display low synonymous (and low protein) diversity. As core genes are not expected to be frequently added to relatively small bacterial core genomes, such sweeps are likely to be interrupted before fixation, a process recently described for SNPs termed adaptive tracking (Song *et al*. 2025). Although we find very limited evidence of this process in our analysis, the lack of power our test has for low-frequency AGs means we cannot conclude adaptive tracking is not a potentially important force in AG evolution.

To address the timescale over which AGs are maintained under balancing selection, we applied a simple calculation. Mean synonymous diversity among the high diversity outlier genes is 0.029 (range across genes 0.002 - 0.095). *K. pneumoniae* is estimated to accumulate mutations at a rate of approximately 6.3 × 10^-7^ per site per year (Moradigaravand *et al*. 2017), which implies that outlier AGs have been maintained for ∼45,000 years (range 3,000 - 150,000 years). This is an order of magnitude deeper than non-outlier AGs (∼3,000 years) and comparable to *θ* estimated from the core genome (∼49,000 years), suggesting that AGs likely under balancing selection have persisted since the species itself diversified. No per generation mutation rate estimate for *K. pneumoniae* is available, but if we use the geometric mean mutation rate from the compilation of mutation rates from 35 species (4.07 × 10^-10^; Lynch *et al*. 2023), we estimate that outlier AGs are approximately 7 × 10^7^ generations old. As an internal check, dividing the accumulation rate by the average mutation rate (Gibson *et al*. 2018), implies that *K. pneumoniae* undergoes ∼1,500 generations per year, or 4.2 per day; corresponding to a doubling time in the wild is ∼5.7 hours, consistent with independent estimates for enteric bacteria.

There are two caveats to our approach. First, recombination allows AGs to depart from the evolutionary trajectory of its background, so if recombination is infrequent, outliers may reflect selection on a linked locus or on the genetic background rather than on the AG itself. Consistent with this, 159 of 171 outliers (93%) that fell within a co-occurrence cluster directly neighboured at least one other gene in that cluster. Second, our filters for multiple introduction might not have excluded genes that are frequently acquired at site-specific recombination hotspots, inflating synonymous diversity of AGs, as such sites hold a large share of accessory gene families (Oliveira *et al*. 2017). However, only genes at a hotspot that arrived from closely related donors would survive both our filtering criteria, so we expect this class to be only a small proportion of the outliers.

Our results demonstrate the value of a frequency-conditioned coalescent expectation for distinguishing not only the extent but also the mode of selection on AGs. We show that at least 29.6% of testable AGs deviate from neutrality, and that most of these show long-term accumulation of synonymous diversity consistent with balancing selection rather than recent sweeps. Beyond detecting balancing selection, our approach offers a first approximation of its timescale, which suggests that these AGs may have been maintained for much of the species’ history. Extending this framework to incorporate variable mutation and homologous recombination rates, and ecological data, would allow direct tests of the mechanisms maintaining this variation. Long-term balancing selection may thus be a widespread and underappreciated force shaping the accessory gene content of bacterial species.

## METHODS

### Genome collection, sequencing and filtering

*Klebsiella pneumoniae* genomes were obtained from the SpARK One-Health study; sample collection and short-read sequencing procedures are described in (Thorpe *et al*. 2022). Hybrid assemblies combining short- and long-read data (European Nucleotide Archive project numbers: PRJEB27342, Illumina sequences; PRJEB66363, Oxford Nanopore), having previously been assembled using Unicycler v0.4.8 (Wick *et al*. 2017). Genomes that did not circularise during assembly were excluded from analyses. From the 412 hybrid genomes, replicate clonal isolates were excluded via hierarchical clustering performed on pairwise Mash distances (Ondov *et al*. 2019), resulting in 260 complete chromosomal genomes available for analysis. O loci and O types were assigned to the chromosome assemblies using *Kaptive* (v3.3.2; Stanton *et al*. 2025) with the *K. pneumoniae* species complex O database (v3.2.1; Whitfield *et al*. 2025).

### Pangenome construction

Genomes were annotated with Bakta (Schwengers *et al*. 2021) and the pangenome was constructed using PIRATE (Bayliss *et al*. 2019) using identity cutoffs of 95–100% in 1% increments and CD-Hit lowest percent identity of 100%, and excluding HSPs below 60% of the aligned sequence length. PIRATE was run with the paralog identification command; after accounting for fission and fusion events, genes were considered paralogs when average gene copy number was greater than 1 (dosage >1). Fission and fusion alleles were excluded from alignments due to poor alignment quality, though their gene families were retained. Genes were considered core if present in more than 257 isolates (>99%). Gene-specific multiple sequence alignments were generated with MAFFT v. 7.310 (Katoh *et al*. 2002). When reporting gene names, to avoid over-attributing function, a gene symbol is used only where it appears within the annotated product, and the product is used in all other cases.

### Per-gene diversity and segregating site estimation

Synonymous sites were identified within each gene alignment using third-position degeneracy. Using custom R scripts, nucleotide counts at the third position were recorded per sequence, with the effective sample size adjusted to exclude sequences carrying missing or ambiguous bases (’N’) at that codon. Throughout, four- and two-fold degenerate sites were treated independently, with estimates reported as the unweighted mean across both classes. For all gene alignments, we recorded the alignment length and the number of synonymous sites. For core genes, both Watterson’s *θ* and synonymous nucleotide diversity, π_S_, were estimated for 2*N_e_μ* to scale coalescent trees.

### Synteny analysis using PanGraph

We chose not to try to infer multiple introductions from the phylogeny (i.e. a single introduction should appear consistently across a clade) as recombination is prevalent in bacteria such as *Klebsiella pneumoniae*. Syntenic blocks were identified using the PanGraph program (Noll *et al*. 2023), where genomes were aligned with minimap2 (Li 2018), setting the alignment sensitivity to accommodate 5% divergence. The fragmentation penalty α and minimum block length were both set to 500 to prioritise the recovery of large syntenic regions suitable for gene-order analysis, suppressing short blocks arising from minor structural variants. Core regions ≥500 *bp* identified by PanGraph were grouped into consecutively oriented ’core-blocks’ using *pypangraph.py*. This produced 30 collinear segments of the core genome, whose endpoints coincide with sites of structural divergence among the isolates analysed. Independent introduction events were inferred when AGs were localised to different core collinear blocks or to distinct genomic regions within core collinear blocks.

### Phylogeny

A core genome alignment was generated from the PanGraph output using SPARK_1006_C1 as the reference strain and processed with Gubbins (Croucher *et al*. 2015) to remove regions of homologous recombination. The resulting recombination-filtered alignment was used to construct a phylogenetic tree using FastTree (Price *et al*. 2010) for the initial tree construction followed by 10 iterations of RAxML (v8, Stamatakis 2014) with 100 bootstraps.

### Identifying mobile genetic elements

Insertion sequences and transposons were identified using both the IS element detection via ISFinder from the Bakta v1.8.1 annotation and complemented with ISEScan (Xie and Tang 2017) for improved sensitivity. Integrons were found using IntegronFinder (Néron *et al*. 2022) and prophages with PHASTEST (Wishart *et al*. 2023). Integrative and conjugative elements (ICEs) and integrative and mobilisable elements (IMEs) were predicted with ICEfinder (v2.0; Wang *et al*. 2024).

### Simulating the neutral distribution of accessory genes

To derive the neutral expectation for the relationship between the number of synonymous segregating sites and AG frequency, assuming a constant effective population size, we modelled the genealogy of the AG insertion site under a standard coalescent model (Hudson 1990). Genealogies of 260 haploid genomes, without recombination, were simulated using *msprime* (v1.4.2, Baumdicker *et al*. 2022). For the *i*th genealogy and each frequency class *k*, where *k* is the number of genomes in which the AG was observed, we identified the sub-genealogy of the *k* descendants. The branch lengths of these sub-genealogies were summed to yield *t_ik_* (Fu 1995). In total, 500,000 independent coalescent trees were simulated, as variance of the largest subtree was found to stabilise at this number of replicates (Fig S3).

Branch lengths, in units of *N_e_* generations, were scaled by *θ/2* to give the number of expected synonymous segregating sites. Because the number of synonymous sites differs among genes, this expectation was evaluated separately for every AG rather than once per frequency class:

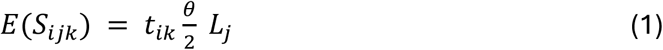

where *L_j_* is the number of synonymous sites in gene *j*, *θ* was nucleotide diversity, *π_S_*, estimated from the core genes (see below for the interpolated θ^, accounting for an expanding population). Then using this expectation, we calculated the probability of observing segregating sites using the Poisson distribution:

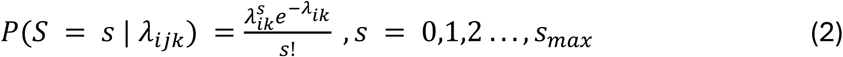

where *λ_ijk_* equals *E*(*S_ijk_*), and *s_max_* was the maximum number of segregating sites across all coalescent replicates. Poisson probabilities were calculated for each sub-genealogy replicate (Tavaré 1984), then summed across replicates and normalised to construct the final expected neutral probability distribution per frequency class. The observed *S*_j_ of each AG was compared against the distribution for its *k* and *L*_j_; two-sided *p*-values were derived using the minimum-tail method, bounded at one. The Benjamini-Hochberg method (Benjamini and Hochberg 1995) was applied to correct for multiple hypothesis testing, using a 5% false discovery rate.

### Calibrating population growth and θ

The expected number of synonymous segregating sites in an AG depends on the population-scaled mutation rate, *θ* = 2*N*_e_*μ*, and on the shape of the genealogy. We estimated both from core genes (present in ≥99% of genomes). Using *msprime* (v1.4.2, Baumdicker *et al*. 2022), we simulated genealogies under population expansion. As in the constant size model, we sampled 260 haploid lineages (ploidy = 1) from a single panmictic, non-recombining population. Setting present day size of 1, places time in coalescent units. Expansion was modelled as deterministic exponential growth at rate α, giving *N*_(t)_ = *N*_0_e^(-αt)^ backwards in time; where α = 0 is a constant size. As *π_S_*/*θ*_W_ is independent of *θ* and depends only on the growth rate, *α*, and sample size, we simulated 10,000 genealogies across a range of *α* between 0 and 500 and interpolated the expected ratio, on a log_10_(1 + *α*) scale. Fixing *α*, we then simulated 10,000 genealogies under the infinite-sites model at each value of *θ* from 0 to 0.1 in steps of 0.005 and interpolated the expected *π_S_* to the observed value.

### Functional analysis for outliers

AGs that were identified as outliers were functionally annotated using eggNog-mapper (v 2.1.13, Cantalapiedra *et al*. 2021) and eggNOG database (v7, Hernández-Plaza *et al*. 2026), augmenting and updating the COG assignments produced by Bakta, which internally calls an earlier version of eggNOG database. eggNOG-mapper was run on the AG representative sequences from PIRATE gene clusters, using DIAMOND (v2.0.15; Camacho *et al*. 2009; Buchfink *et al*. 2015) with the following parameters: sensitive mode; a minimum alignment score of 60; a minimum percentage identity of 40%; query and subject coverage thresholds of 20%, sufficient to capture partial matches to conserved functional domains in assembly fragmented genes; automatic taxonomic scoping; and, inclusion of all ortholog types. The resulting COG ids were mapped to the COG 2024 database release (Galperin *et al*. 2025). To test whether outlier AGs were enriched for particular COG functional categories, Fisher’s exact tests were performed comparing category membership between outlier and non-outlier AGs, excluding genes lacking COG assignments, with Benjamini-Hochberg correction (Benjamini and Hochberg 1995).

### Protein allelic diversity

Protein allelic diversity was calculated for each accessory gene as Nei (1978)’s unbiased gene diversity, *H* = (*N*^2^ – ∑*n*^2^_*i*_)/(*N*(*N* – 1)), where *N* is the number of genomes carrying the gene and *n*_i_ the number carrying the amino acid variant *i*. *H* is the probability that two of those genomes, drawn at random, encode different proteins.

### Detecting co-occurring genes

Goldfinder (Gavriilidou *et al*. 2024), a phylogeny aware gene association tool, was used to detect co-occurrence and avoidance between AGs. The input phylogeny was the recombination-corrected core-genome tree produced by Gubbins from the PanGraph syntenic core alignment. Using the default settings, AGs present in fewer than 5% of isolates were excluded. Both associations and dissociations were tested using the scoring methods: simultaneous and subsequent (Collins and Didelot 2018). The terminal and Coinfinder scoring methods were also evaluated but yielded no significant hits. Multiple testing was accounted for using false discovery rate correction (α = 0.05). A Fisher’s exact test was performed to assess whether outlier AGs were enriched among all significant Goldfinder associations (co-occurrence and avoidance combined).

### Detecting ecological association genes

We implemented treeWAS to detect genes that were associated with isolation sources while accounting for phylogenetic structure (Collins and Didelot 2018). As in our Goldfinder analysis, the gene presence/absence matrix was restricted to intermediate-frequency genes (>5%, i.e., present in >12 of 260 genomes, excluding fixed genes) and tested against binary isolate categories on the Gubbins recombination-corrected phylogeny. We used the terminal, simultaneous and subsequent association scores with FDR correction (*q* < 0.05). To ensure both contrasts had sufficient isolates within each category, we examined two categories: hospital (*N* = 167) vs non-hospital association (*N* = 93) and, within human isolates, disease (*N* = 95) vs carriage (*N* = 76). Ecological source, (human, animal or environmental) and hospital association were near-redundant (Cramér’s V = 0.97); we therefore tested hospital association alone.

## DATA AND RESOURCE AVAILABILITY

All code used in data analyses and visualisation is available in GitHub at: https://github.com/CaraConradsen/kpne_ags. The sequencing dataset supporting the conclusions of this article was generated by the SpARK One-Health study (Thorpe et al., 2022) and is available in the European Nucleotide Archive (ENA) under the accession numbers PRJEB27342 (Illumina short-read sequences) and PRJEB66363 (Oxford Nanopore long-read sequences). Accession numbers for the 260 Illumina short-read genomes analysed here are listed in Supplementary Table 2.

## AUTHOR CONTRIBUTIONS

A.E.W., C.C., E.F., M.V. and S.B. conceived and designed the study. A.E.W. designed the sampling scheme. M.V. and A.E.W. developed the test for the accumulation of synonymous segregating sites in accessory genes. C.C. performed the analyses. C.C. and M.V. wrote the manuscript, with contributions from all authors.

## Supporting information

supplementary paper 1

## ACKNOWLEDGEMENTS

This study was supported by funding from the Leverhulme Trust (grant number RPG-2024-212). We acknowledge the use of the University of Exeter’s Advanced Research Computing facilities and thank the ARC team for technical support. We thank Jukka Corander for providing SPARK data.

## Notes

### Competing Interest Statement

The authors have declared no competing interest.

