## supplementary information for "Evidence of widespread balancing selection in a bacterial pangenome"

### Supplementary Information: Evidence of widespread balancing selection in a bacterial pangenome

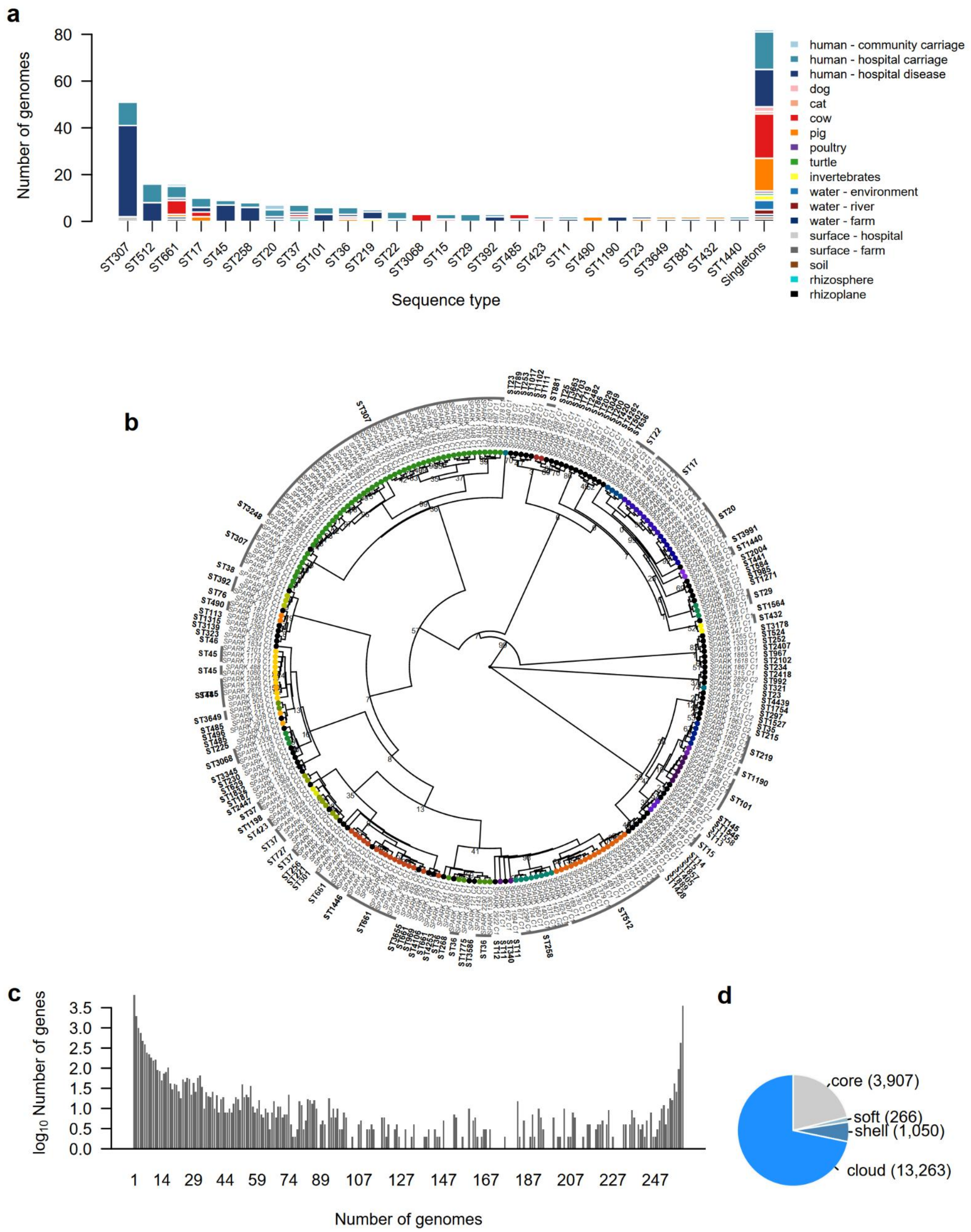


**Figure S1 Isolation source, phylogeny and pangenome composition of 260 hybrid SpARK *Klebsiella pneumoniae* genomes.** (a) Distribution of sequence types (STs) and isolation sources. Bars show the number of genomes per ST, stacked and coloured by isolation source (human, animal and environmental). STs represented by a single genome are pooled as singletons. (b) Maximum-likelihood circular phylogenetic tree of *K. pneumoniae* based on core genes, generated using *RAxML*. Bootstrapped using 100 replicates, where nodes with <100 replicates indicated on tree. Coloured dots specify STs with multiple isolates, while black denotes STs with only one isolate. Inner tips are labelled with isolate names; outer labels correspond to STs, with bars indicating groups. (c) Gene frequency distribution across the *K. pneumoniae* pangenome, where counts are log-transformed to improve resolution. (d) Counts of core genes (present in ≥99% of genomes), soft-core genes (≥95% and <99%), shell genes (>15% and <95%), and cloud genes (≤15%), following Douglas and Shapiro (2021), adapted from Page *et al.* (2015). The number of genes (in brackets) per category are shown.

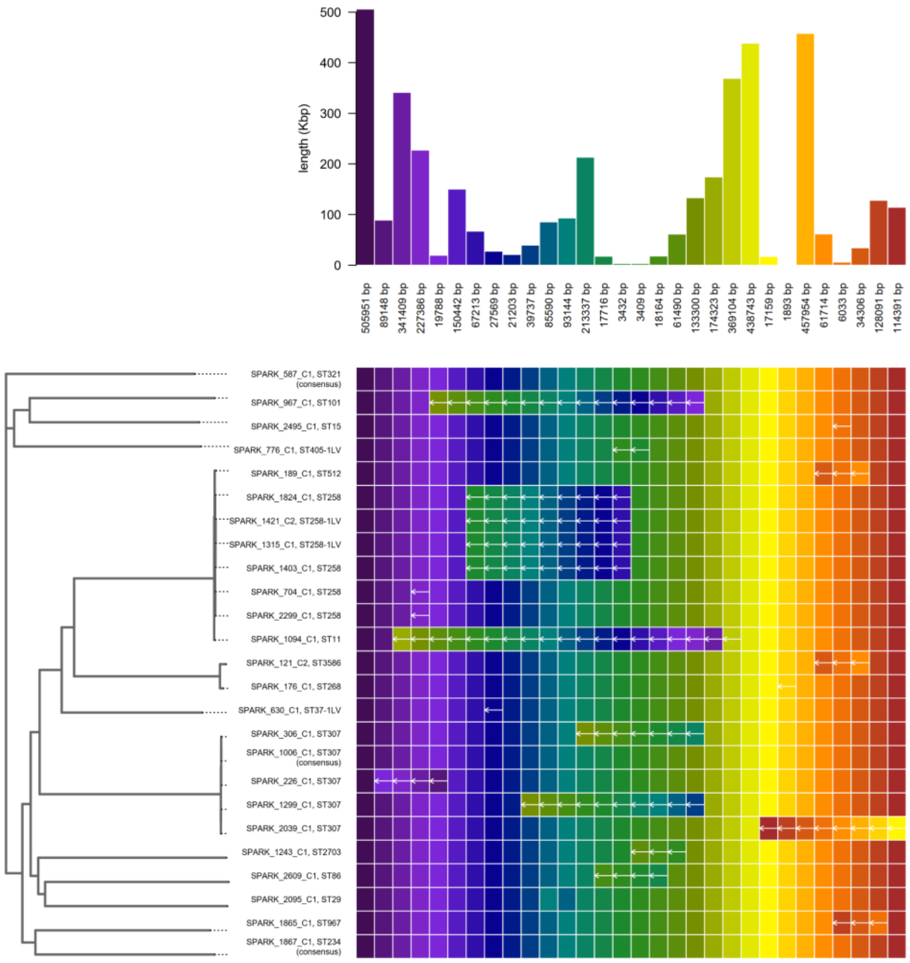


**Figure S2 Structural variation in core genome synteny among *Klebsiella pneumoniae* isolates.** Core regions that were conserved in both orientation and order across all genomes were defined as core syntenic blocks, represented here as coloured blocks. Core syntenic blocks were identified using PanGraph (Noll *et al.* 2023) and grouped using *pypangraph.py*. Rows represent genomes (genome id, sequence type), with only a small number of isolates (22 out of 260) deviating from the consensus order (exemplified by isolates SPARK_587_C1, SPARK_1006_C1, SPARK_1867_C1). White arrows indicate genomic inversions, and changes in coloured block order indicate translocations. The top panel shows a bar graph of block lengths; the left panel shows the recombination filtered core genome phylogeny. The largest inversion is seen in the ST11 genome, SPARK_1094_C1; interestingly, the only other ST11 genome does not have this inversion, retaining the consensus order. A large (9 blocks) shared inversion was detected in four of eight ST258-related isolates.

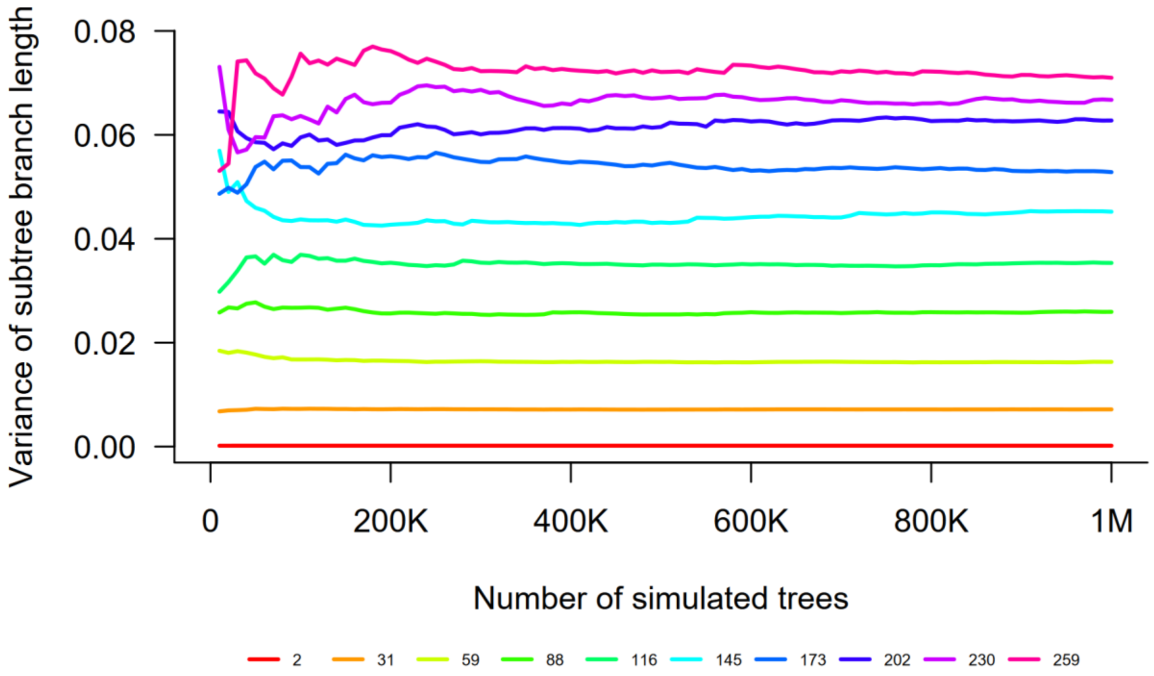


**Figure S3 Variance in branch-length estimates by frequency class k.** Estimates were obtained from 1,000,000 independent coalescent genealogies simulated under a constant population size. For each sub-genealogy relating *k* descendant lineages, branch lengths were summed and the variance in this sum estimated across replicates. Each coloured line represents a single frequency class *k*, from *k* = 2 to 259.

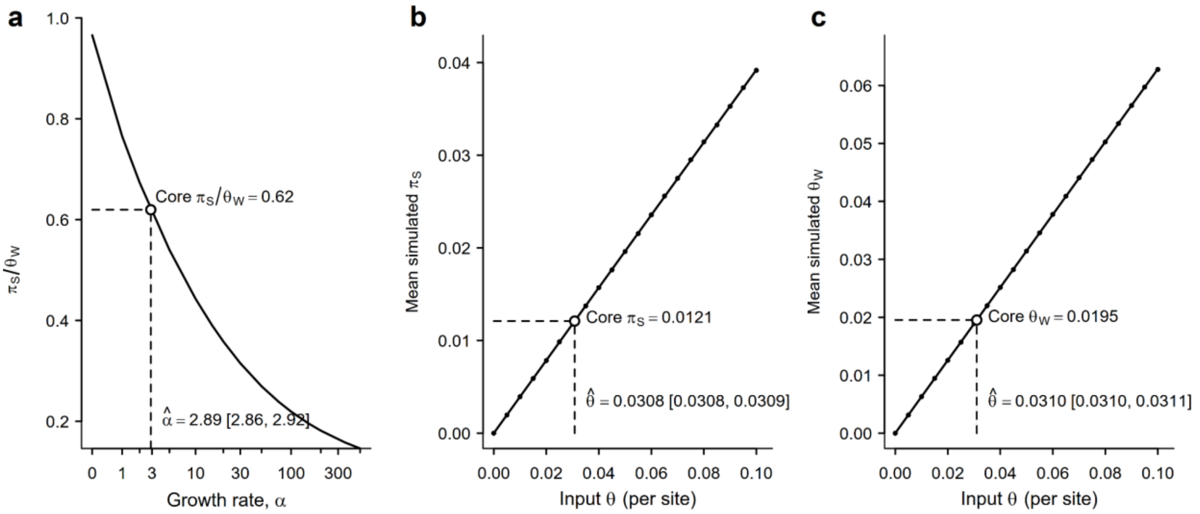


**Figure S4 Two-step interpolation to infer coalescent null expansion parameters using core gene synonymous diversity.** (a) Expected $\pi_S$/ $\theta_W$ as a function of the growth rate, where α is identified without reference to the mutation rate, θ; the x-axis is spaced on $\log_{10}$ (1 + α). Mean simulated $\pi_S$ (b) and $\theta_W$ (c) as a function of the input θ, with growth rate fixed at $\hat{\alpha}$ estimated in (a). For each grid input value, 10,000 genealogies were simulated in *msprime* (Baumdicker *et al.* 2022), and expected values estimated. Confidence intervals were obtained by resampling the simulated replicates within each grid point and re-solving the interpolation, reporting 95% percentile intervals over 1,000 resamples.

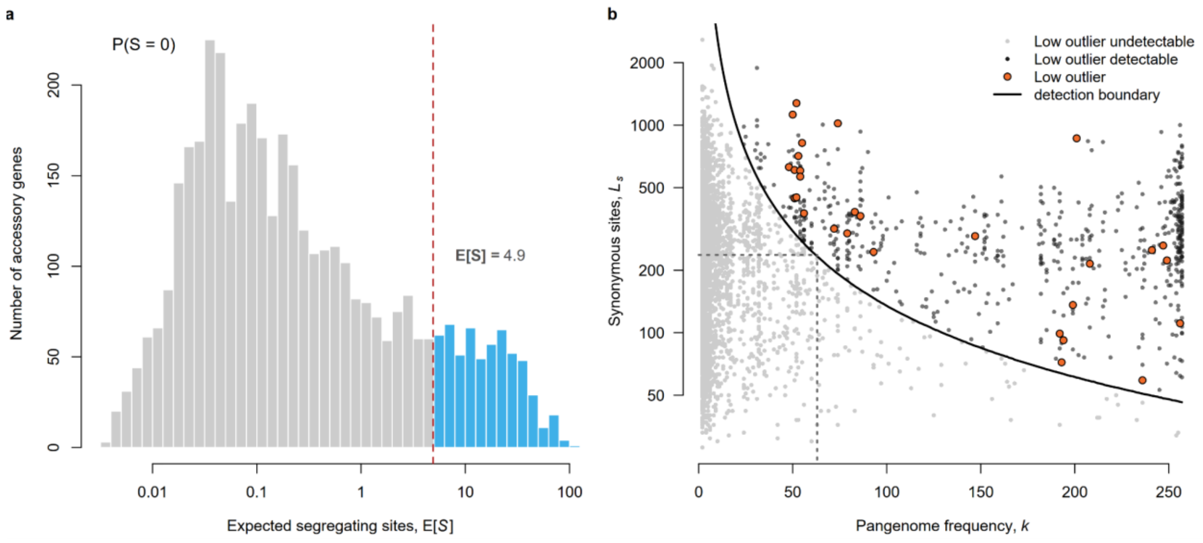


**Figure S5 Detection limits for outliers with low synonymous diversity.** (a) Distribution of the expected number of synonymous segregating sites ($E[S]$), across the 4,071 accessory genes. The most extreme low diversity observation is when $S = 0$ [P($S = 0$)], making the smallest two-sided $P$ value a gene can then return as twice $e^{-E[S]}$. Using the realised Benjamini–Hochberg (Benjamini and Hochberg 1995) critical value ($P = 0.0146$ at $q < 0.05$) places the limit at $E[S] = 4.9$ (red dashed line). Below this value (grey bars), no observed value of $S$ can be determined as a significant departure versus above this value (blue bars). (b) The same limit as in (a) but in the context of pangenome frequency, $k$, and synonymous site count, $L_S$. Since $E[S]$ depends on the product of $k$ and $L_S$, the detection limit for low outliers is curved (black line). Points are individual accessory genes, coloured by whether a low outlier is undetectable (light grey) or detectable (dark grey). Orange points represent the low outliers detected in this study. Dashed lines demarcate the median synonymous length ($L_S = 237$; horizontal), which first becomes testable at $k = 63$ (vertical).

##### Co-occurrence and ecological association of accessory genes under selection

The non-random association of functionally-linked AGs is a well-established feature of bacterial pangenomes (Rosconi *et al.* 2022). Patterns of AG co-occurrence and co-avoidance can be interpreted as signatures of epistatic selection and have been used as evidence that the accessory genome is maintained by selection rather than neutral processes (Goyal 2018; Whelan *et al.* 2021; Beavan *et al.* 2024). We employed Goldfinder (Gavriilidou *et al.* 2024) to identify AGs that co-occur or avoid one another more often than expected given their shared ancestry, and asked whether selection outliers were over-represented. Goldfinder only tests AGs present in at least 5% of genomes ($k \geq 13$), where 428 of the 1,204 outliers (398 with excess and all 30 with deficit diversity) could be assessed against 831 non-outliers. Of these outliers, 171 (40%) were involved in at least one significant co-occurrence association (Supplementary Fig. 6). Relative to non-outliers, associations were more frequent among balancing selection candidates (161 of 398, 40.5%; OR = 1.31, $P = 0.036$) but not among directional selection candidates (10 of 30; OR = 0.96). However, neither comparison survived correction. Of the 171 outliers, 159 (93%) shared a syntenic region with at least one other gene in the same association group, and 144 (84%) AGs were in clusters that all localised to the same region, suggesting that much of this signal reflects co-transfer of physically linked genes rather than epistasis between unlinked loci.

The SPARK dataset records hospital vs non-hospital association origin of isolates, allowing us to ask whether selection outliers are distributed non-randomly between these two sources. We tested the two best-sampled contrasts: hospital-associated ($N = 167$) versus non-hospital ($N = 93$) isolates, and, among human isolates, disease ($N = 95$) versus carriage ($N = 76$) (Supplementary Figure 1a). Ecological associations were tested in the same framework as gene co-occurrence, implemented using treeWAS (Collins and Didelot 2018). Neither contrast linked isolation source to selection candidates: hospital vs non-hospital association recovered five significant genes, none of which was an outlier; however, two genes did belong to a single co-occurrence cluster (57 genes) containing two high diversity outliers, and disease vs carriage recovered none. Most hospital isolates derived from a single hospital (118/167), strongly confounding the source with clonal lineage and so this negative result may reflect either no source-associated selection or limited power to separate ecological from phylogenetic signal.

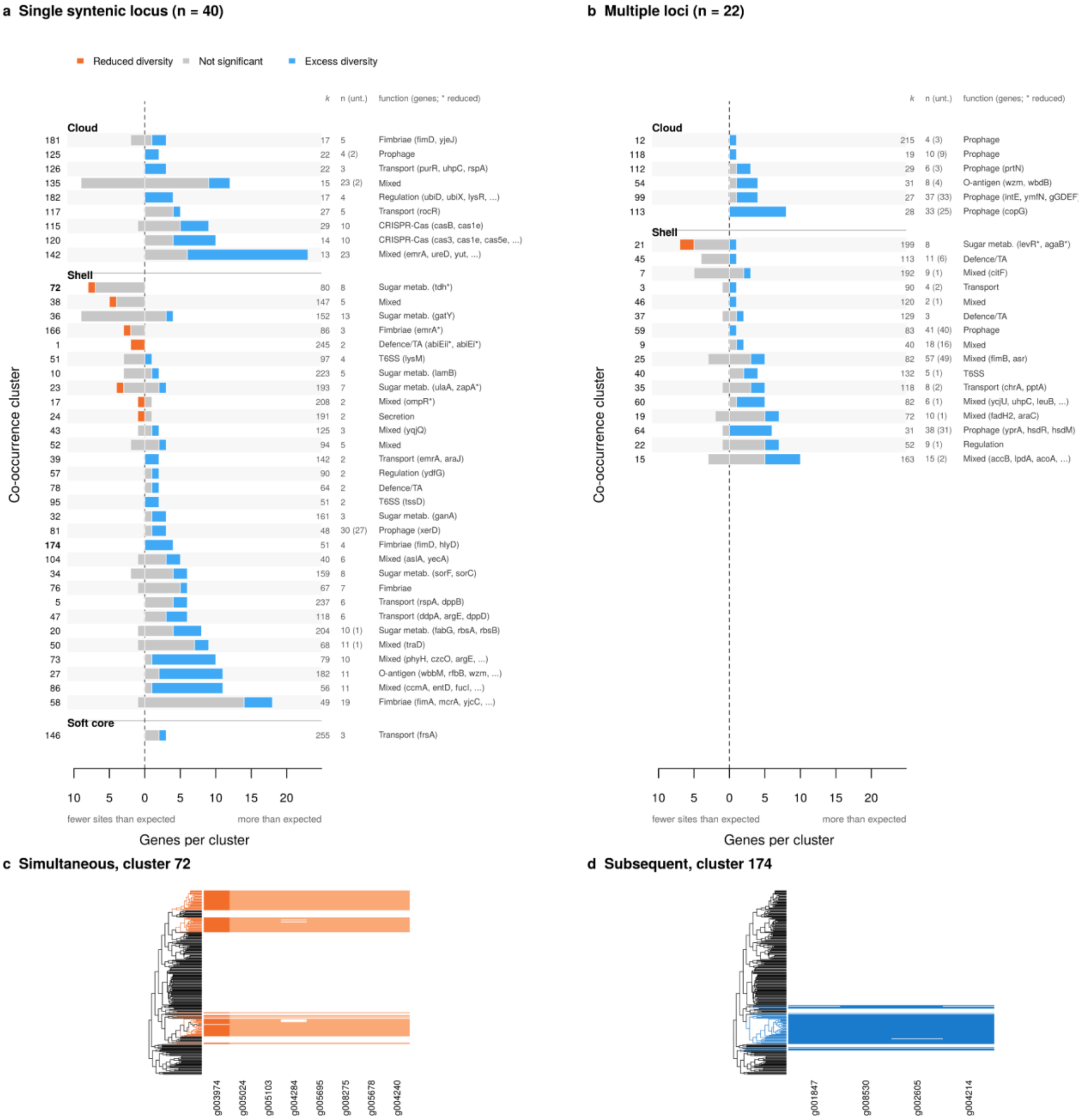


**Figure S6 Co-occurrence clusters containing selection outliers in *Klebsiella pneumoniae*.** Goldfinder co-occurrence analysis (Gavriilidou *et al.* 2024) identified genes co-occurring beyond phylogenetic expectation, scored as simultaneous (coincident gain/loss) or subsequent (sequential acquisition, then maintenance). Goldfinder imposes a minimum 5% frequency threshold for hypothesis testing; co-occurrence among outliers present in fewer than 13 genomes therefore cannot be detected. (a, b) Distribution of synonymous diversity signals within clusters containing at least one outlier; separated into clusters whose members all occupy a single syntenic locus (a; $N = 40$) and clusters whose members sit at multiple loci (b; $N = 22$). Each row is a cluster, and each member gene is counted on the side of the neutral expectation it falls on, with fewer synonymous sites than expected on the left and more than expected on the right. Genes are coloured by deviations of synonymous diversity against the neutral coalescent model (orange, low diversity; light grey, not significantly different from neutral expectation; blue, high diversity). Clusters are grouped by pangenome class and ordered within each class from rows favouring reduced diversity to those favouring excess diversity. Margins give the cluster median pangenome frequency ($k$), the number of member genes with the number lacking a neutrality test in parentheses, and the inferred function with named outlier members. (c, d) Examples of presence/absence of cluster recovered by the simultaneous score (c; cluster 72, eight genes) and by the subsequent score (d; cluster 174, four genes), mapped against the Gubbins recombination-corrected phylogeny (tree distance on a log scale). Filled bars denote presence; in both clusters all member genes co-occur invariantly.

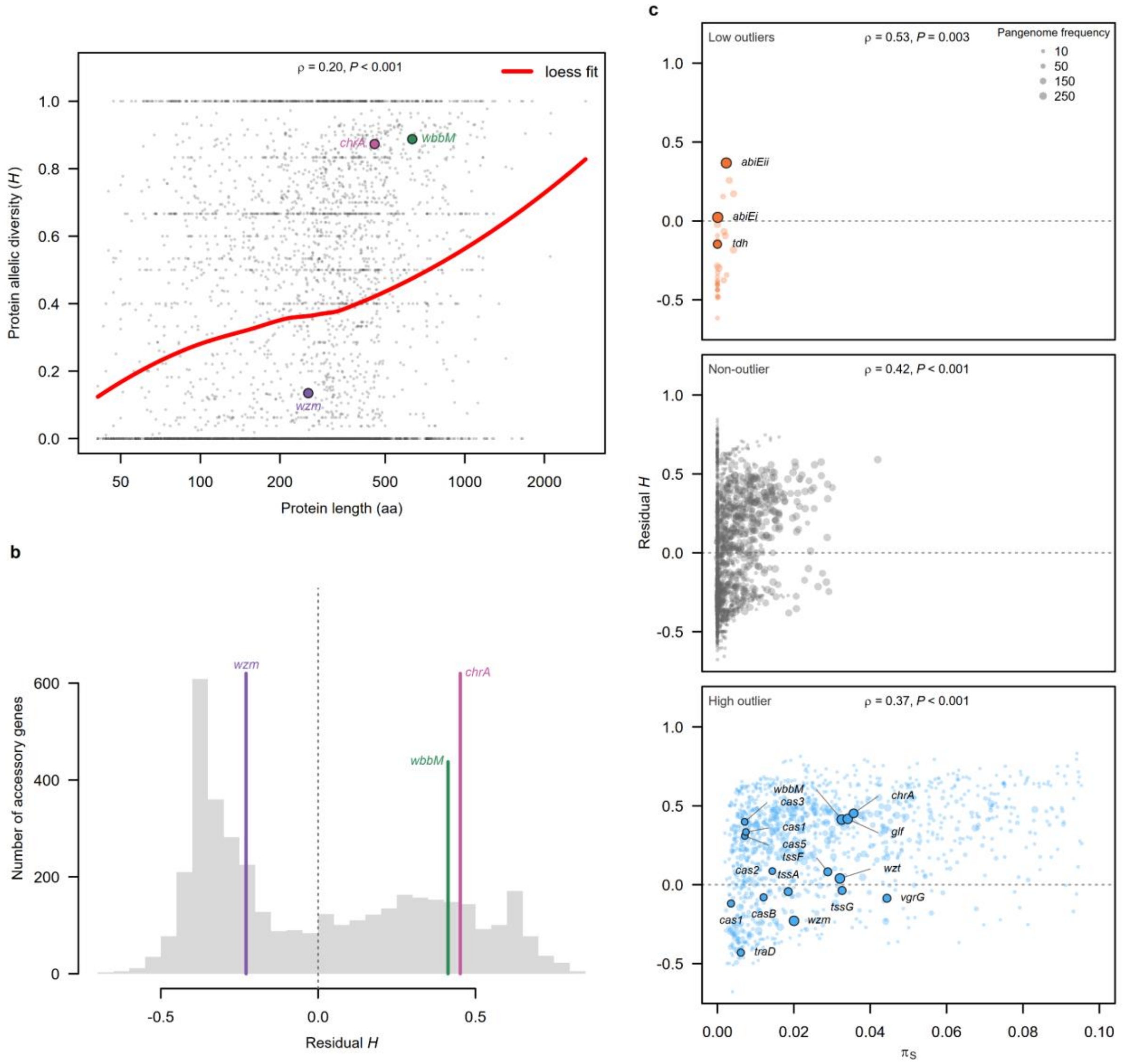


**Figure S7 Protein diversity corrected for gene length.** (a) Nei (1978)'s unbiased protein diversity, *H*, against protein length (number of amino acids, log scaled). A loess fit of *H* against log gene length (smoothing parameter of 0.75, red line), indicates that gene length accounts for 4.8% of the variance in *H*. Residuals from this fit were used to correct for gene length in the following panels, b and c. Spearman's rank correlation, *ρ*, between *H* and protein length is given at the top of the box. *wzm, wbbM* and *chrA* (Fig. 4b–d) are marked. (b) Distribution of residual *H* across accessory genes (AGs). The dotted vertical line marks zero and coloured lines mark focal AGs: *wzm, wbbM* and *chrA.* (c) As in Fig. 4a (Main text: Results), with residual *H* instead of *H.* Here, the horizontal dotted line marks zero.

**Table S1 Accessory genes (AGs) with significant departures from neutral expectation.** Outliers (*N* = 1,204; 1,174 high, 30 low) were classed as high [low] diversity outliers if there were more [fewer] synonymous segregating sites than expected under neutrality (Outlier type). Gene families (Gene) and pangenome frequency (Gene freq.; number of genomes carrying the gene) were determined using PIRATE (Bayliss *et al.* 2019), where AGs are classed as soft-core, shell or cloud (present in ≥95% and <99%, >15% and <95%, and ≤15% of genomes, respectively). Synonymous nucleotide diversity (*pi_s*) estimated across fourfold and twofold degenerate sites, Synonymous sites is the number of these sites assayed, Segregating synonymous sites (*S*) is the observed count, and *E*[*S*] is the number expected under the neutral coalescent null. Gene length (*bp*) is that of the longest ortholog in each cluster. The *P*-value and the Benjamini-Hochberg 5% false discovery rate-corrected *P*-value (FDR P; Benjamini and Hochberg 1995) are reported. COG category provides the Clusters of Orthologous Groups functional classification (Cantalapiedra *et al.* 2021). Tidy gene name and Consensus product were annotated using Bakta (Schwengers *et al.* 2021). Phage Completeness and Phage Type give the number of genomes in which the gene lies within a PHASTEST prophage region, by completeness, and the most common prophage hit (Wishart *et al.* 2023). ICE and Insertion sequence Family give the number of genomes in which the gene lies within an ICEfinder-predicted integrative and conjugative element (Wang *et al.* 2024) or an ISEScan insertion sequence family (Xie and Tang 2017). Core-block number and Flanking core genes give the syntenic core block containing the gene and the core genes flanking its insertion site. Co-occurrence group and Detection method give the Goldfinder co-occurrence cluster and scoring method (Gavriilidou *et al.* 2024).

**Table S2. *Klebsiella pneumoniae* genomes analysed.** The 260 genomes were part of the SpARK project (Thorpe *et al.* 2022). Accessions are for the genome data in the European Nucleotide Archive (project PRJEB27342). Sequence types (STs) are from Thorpe et al. (2022), with single- and multi-locus variants assigned to their parent ST. O loci and O types were assigned to the chromosome assemblies with *Kaptive* (v3.3.2; Stanton *et al.* 2025) using *K. pneumoniae* species complex O database (v3.2.1; Whitfield *et al.* 2025), genomes *Kaptive* classed as untypeable are labelled as such.
